# Combined Raman Microscopy and Transmission Electron Microscopy shows the co-existence of whitlockite crystals and carbonated hydroxyapatite-mineralized collagen fibrils in human calcified aortic valves

**DOI:** 10.1101/2025.03.03.641198

**Authors:** R.H.M. Van der Meijden, C.T.A. Kuster, A. van Broekhoven, R. Allgayer, L Rutten, R. Roverts, M. Cerruti, E. Macías-Sánchez, J.H. Cornel, N. Sommerdijk, S. El Messaoudi, A. Akiva

## Abstract

Aortic valve stenosis (AS), the leading valvular disease in aging populations, is driven by complex extracellular matrix (ECM) modifications and progressive calcification. Despite its clinical significance, the molecular mechanisms underlying AS remain poorly understood, limiting therapeutic options to invasive valve replacement. This study employs Raman microscopy, a label-free imaging technique, and its integration with transmission electron microscopy (TEM) to elucidate the biochemical and ultrastructural changes in stenotic aortic valves. High-resolution spectroscopic imaging revealed distinct extracellular matrix modification across different regions, involving elastin degradation, cholesterol deposition, and mineral formation, comprising not only hydroxyapatite (cHAp) but also whitlockite.

Elastin-rich domains associated with cHAp deposition exhibited cross-link degradation, while collagen matrices supported mineralized phases with varying mineral-to-matrix ratios that, in heavily mineralized regions, went far above those of mature human bone. For the first time we demonstrated whitlockite as a mineral deposit in calcified aortic valves, in areas with different degrees of calcification. This implies that this Mg containing mineral, which has been considered a precursor to cHAp in pathological calcification, forms independently, challenging prevailing models of calcification. The combination of Raman and TEM showed how bone-like cHAp mineralized collagen matrix in later stages engulfs the initial non-physiological whitlockite deposits.

This correlative multimodal approach advances our understanding of AS by capturing spatially resolved chemical and structural dynamics at the nanoscale. The findings highlight Raman microscopy’s potential for probing calcification mechanisms across diverse tissue types and suggest its role in identifying novel therapeutic targets. This study underscores the value of integrative imaging methodologies in unraveling complex pathological processes and advancing patient care.

## Introduction

In the developed world, one in four people aged >65 years suffer from calcific aortic valve disease (CAVD), which is the underlying cause of aortic valve sclerosis and subsequent aortic valve stenosis.[1] Aortic valve sclerosis is characterized by the thickening and calcification of the heart valve; in 10-15% of cases, this progresses to aortic valve stenosis (AS).[2, 3] AS leads to obstruction of left ventricular outflow and can cause angina, syncope, heart failure, and sudden cardiac death; with the necessity of medical intervention in the later stages of disease.[4] Once symptomatic, untreated elderly patients have a poor prognosis with a 2- and 5-year survival rate of 40% and 22%, respectively.[5] AS has a major impact on health care and is expected to increase in the coming decades due to the aging population.[2, 6] Currently, aside from invasive valve replacement, there are no available treatment options.[3, 7] A deeper understanding of the disease and its progression could provide better treatment options for patients in the future.

In healthy individuals, the aortic valve consists of three leaflets, each around 500 µm in thickness,[9] which open and close to direct the flow of blood (Fig. 1a). These leaflets consist of a three-layered extracellular matrix (ECM) that is covered on each side by a monolayer of valve endothelial cells (VECs) (Fig. 1b).[10] The composition of the three ECM layers is optimized to provide strength and flexibility to the valve, with the aortic-side layer (fibrosa) rich in collagen, the central layer (spongiosa) in glycosaminoglycans and the ventricular-side layer (ventricularis) in elastin.[11] These ECM layers also house valve interstitial cells (VICs), which are responsible for maintaining the structure and composition of heart valve ECM.[11]

**Fig 1.**
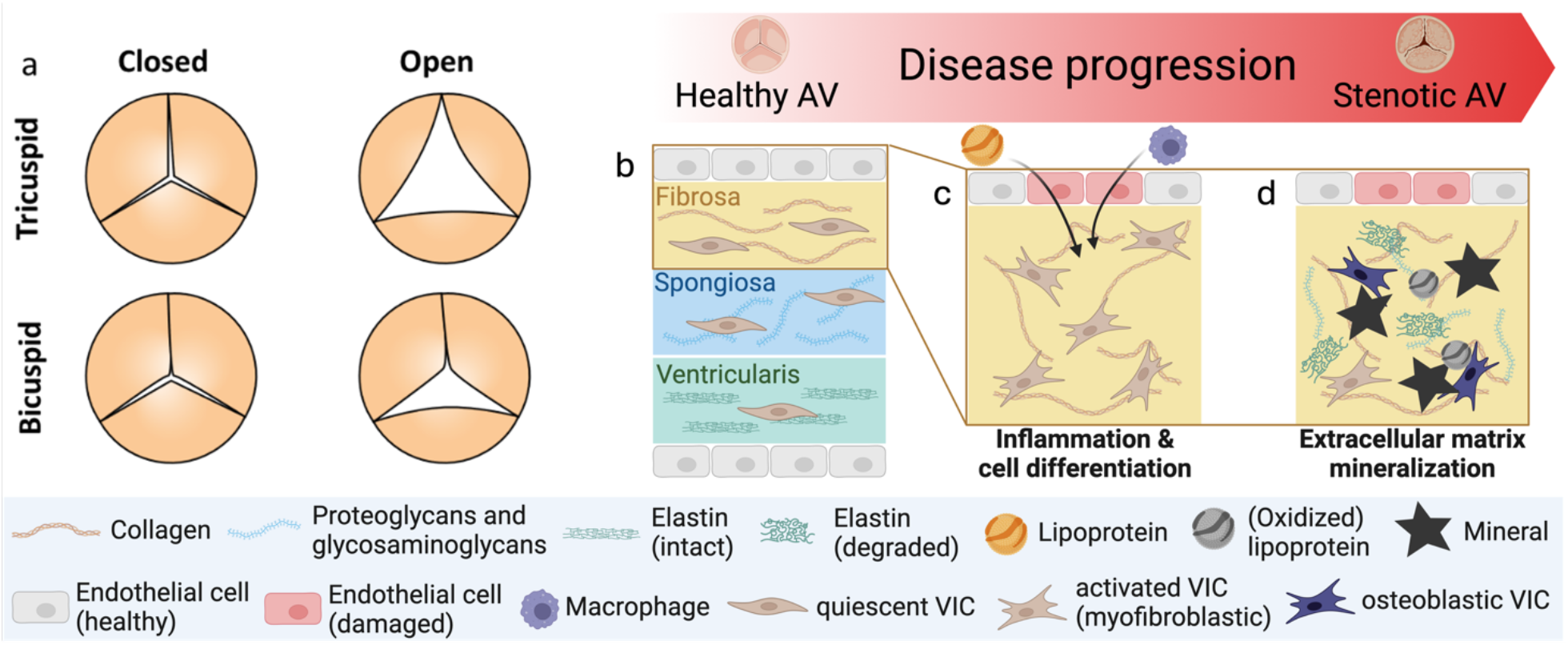
Schematic of the aortic valve. a) Illustration of a tricuspid (top) and one form of bicuspid (bottom) valve during the opening and closing of the aortic valve. Schematic representation of the microanatomy of b) a healthy valve, and c) early and d) late stages of disease progression of CAVD starting from a normal leaflet, followed by infiltration of lipoproteins and immune cells after endothelial damage, resulting in a mineralized valve ECM. VIC: valvular interstitial cell. Image generated using BioRender.[8]

The current proposed pathophysiological process of CAVD starts by damage and stimulation of the VECs,[12] initiated by mechanical[13] or oxidative[14] stress (Fig. 1c). This allows infiltration of oxidized lipoproteins[15, 16] and immune cells[17], creating an inflammatory environment where (1) VECs undergo an endothelial-to-mesenchymal transition into VICs,[18] (2) VICs differentiate into myofibroblast-like and osteoblast-like cells,[19] and (3) a high concentration of macrophages is present.[20] The lipoproteins are enzymatically degraded, resulting in free fatty acids and lysophosphatidylcholine, promoting VIC apoptosis.[21] VIC differentiation towards osteoblastic or myofibroblastic lineages[22] results in an increase in collagen production, leading to valve thickening and subsequential mineralization,[23] which, in return, modify the mechanical properties of the valve (Fig 1d).[24, 25]

Although evidence for partial differentiation of valvular cells towards osteoblast-like cells suggests a parallel to bone development,^[26]^ there are also reports of significant differences in matrix development compared to bone.^[27-29]^ Currently, however, our limited capability to characterize matrix development prevents us from proposing a detailed mechanism for matrix calcification in CAVD. Specifically, there are still many questions on how the disease modifies the organic matrix, how this leads to calcification, and how the mineral develops within the matrix.

By spatially resolved studying the material chemistry underlying the mechanisms of CAVD we can begin answering these questions. This cannot be done *in vivo*, as current clinical imaging strategies do not provide the resolution nor the chemical information to identify the different stages of pathological calcification[30]. However, detailed *ex vivo* analyses of extracted patient materials provide useful insights into this heterogeneous process.[29, 31-33]

Optical microscopy of histologically stained samples is the most common approach for AS tissue analysis. It provides useful biochemical and structural information on the macroscopic level. However, due to the extensive sample preparation required, which includes tissue decalcification, and the limited resolution of optical microscopy, the underlying mechanism cannot be studied in detail using this approach. In contrast, Raman microscopy provides (bio) chemical information without the need for staining while providing semi-quantitative information on both the organic and inorganic components of the matrix.[34-36] Nevertheless, only a handful of studies exploited Raman microscopy for AS investigation[31, 37, 38] and were able to describe details of the composition of the calcification.[31]

Bicuspid aortic valve (BAV) is a congenital anomaly where the aortic valve has two cusps instead of the usual three, affecting approximately 0.5 to 2% of the global population, leading to the narrowing of the valve opening that obstructs blood flow from the heart (Fig. 1a).[39-41] Clinical studies have highlighted that BAV patients face a significantly higher risk of developing AS compared to those with a normal tricuspid aortic valve (TAV).[42] Despite its lower prevalence, BAV is implicated in a notable proportion of AS cases, with research suggesting it may contribute to a substantial burden of AS requiring medical intervention.[42, 43] AS also tends to manifest earlier in life among BAV patients, typically presenting symptoms and requiring medical attention approximately a decade earlier than individuals with a TAV.[43] Even at birth, the ECM of BAVs has often lost its tri-layered structure and the BAV leaflets are subjected to higher hemodynamical stress, both of which could contribute to the higher degrees of mineralization in BAVs.[44] Understanding the mechanisms underlying the clinical factors will be crucial for effective management and treatment strategies in patients with BAV-associated AS.

In the current study, we explore the potential of Raman spectroscopy in exposing the biochemical cues that indicate the progression of CAVD in bicuspid aortic valves. We investigate the composition and structure of the affected valves, quantifying the distribution of the most prevalent components: organic matrix, lipids, and minerals at different locations within a single leaflet. For this, we combined Raman microscopy and transmission electron microscopy (TEM) to obtain semi-quantitative location-specific chemical information together with structural information at nanometer resolution. Our study shows how chemical and structural mapping by Raman microscopy can help to understand the pathways of calcification in CAVD.

## Materials and Methods

### Materials

This study was conducted as part of the “Pathophysology of Aortic Valve Stenosis (Patho-AS)” study at the Radboudumc in Nijmegen, the Netherlands. The use of excised human aortic valves was approved by the Medical Ethics Committee of the Region Arnhem-Nijmegen under file number 2023-16767 and was conducted in accordance with the ethical standards laid down in the 1964 Declaration of Helsinki and its later amendments. All participants gave informed consent prior to inclusion in the study.

Bicuspid aortic valves were collected from patients who underwent elective surgical aortic valve replacement due to severe stenosis or moderate stenosis when there was an indication for coronary artery bypass grafting (CABG). Moderate or severe AS was defined through transthoracic echocardiography according to the 2017 EACVI / ASE guidelines for assessing aortic valve stenosis.[45] For this study, two bicuspid valves with severe stenosis were taken from one male (M-108) and one female (M-178) patient, both 74 years old.

### Experimental

#### Aortic valve sample preparation

A fresh tissue sample from M-108 was selected from the middle of the leaflet spanning its entire transverse direction and embedded in “optimal cutting temperature” (OCT; Tissue-Tek OCT, Sakura FinetekUSA, CA) compound. The sample was then transversally sectioned at -20 °C to 4 μm slices, such that the slices span the entire cross-section of the leaflet. Adjacent sections were subjected to histological staining and Raman spectroscopy imaging, respectively. The sections for Raman analysis were mounted on aluminum foil, rinsed and fixed for 15 minutes in 70% ethanol at room temperature.

To preserve the integrity of the calcified areas, sample M-178 was sliced using a vibratome. For this, tissue sample was fixed in 70% ethanol at room temperature for 24 hours and stored at 4°C before sectioning. Raman samples were prepared by drying the tissue sample and mounting it on a microscopic glass with super glue. The tissue was trimmed using a vibratome (VT 1200S) at a speed of 0.01 mm/s and amplitude of 3 mm, exposing a new internal surface. The mounted tissue sample was then polished to achieve a smooth surface for Raman spectroscopy imaging.

#### Histological Imaging

An OCT cryo-sectioned 4 µm tissue section was stained with Masson’s Trichrome using a standard Trichrome Staining Kit (KK02, Roche, Switzerland).

#### Raman spectroscopic imaging

The WITec alpha 300R confocal Raman microscope system was used for spectra collection. All data were collected using the Zeiss EC Epiplan-Neofluar Disc 50x/0.8 objective, at a lateral resolution of 0.5 µm, and using a 600/mm grating. Raman data of unpolished OCT cryo-sectioned 4 µm tissue sections (Fig. 2) were collected with a 532-nm laser, laser power of 8.0 mW, and 1-s integration time per spectrum. Raman data of the polished vibratome-sectioned tissue sample (Fig. 3) were collected with a 633-nm laser, laser power of 11 mW, and 2-s integration time per spectrum. Here, five regions of 60×60 µm were selected for imaging at 500 nm resolution (Fig 3a). All spectra were collected with a polarized laser.

**Fig 2.**
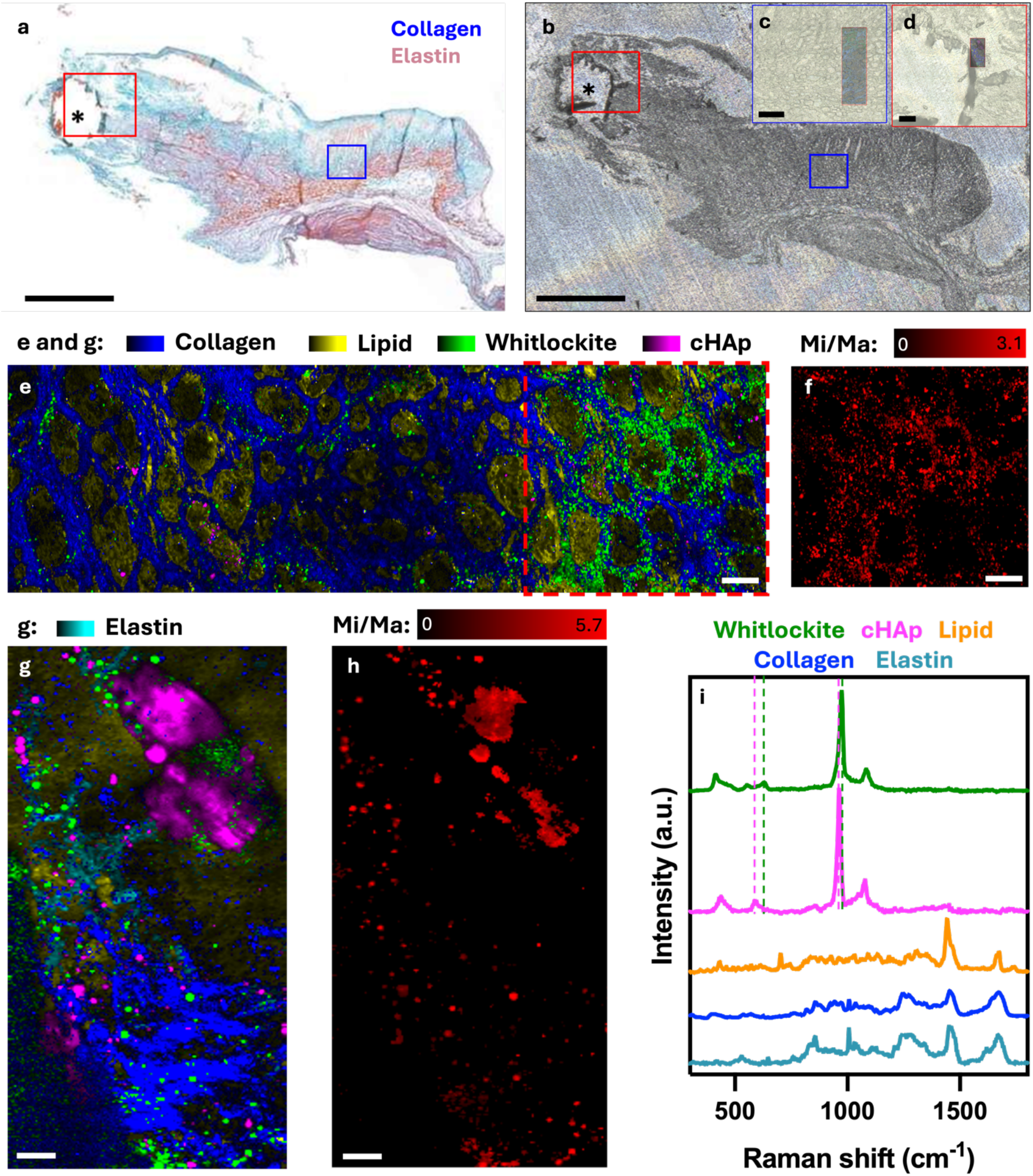
Characterization of cryo-sectioned calcified human aortic valve. a) Overview image of a 4 µm section of the heart valve stained with Masson’s Trichrome staining showing collagen (blue) and elastin (light pink). b) Consecutive section before Raman imaging, placed on an aluminum substrate. The regions of interest for further Raman analysis are marked in blue and red squares in a and b. The location of the visible mineralization is indicated with *. Scale bars: 1mm. c) Overview of the ROI away from the visibly mineralized region (blue box in a and b) Inset: overlay of the Raman image shown in e. d) Overview of the region of interest close to the visible mineralization (red box in a and b). Inset: overlay of the Raman image shown in g. Scale bars: 100 µm. e) Component image based on Raman spectroscopy analysis of the ROI in c. Component analysis highlights four main contributors to the matrix: collagen (blue), lipid (yellow), whitlockite (green) and cHAp (purple). The component map shows the most prominent component in each pixel. The color intensity in each pixel corresponds to the Raman signal intensity. Black areas showed no Raman peaks, i.e. they correspond to holes in the tissue section. f) Overview of the Mi/Ma for the whitlockite-rich ECM region in e, denoted by the dashed square. Scale bars in e and f: 20 µm. g) Component image based on Raman spectroscopy analysis of the ROI in d. Component analysis highlights five main contributors to the matrix: collagen (blue), lipid (yellow), whitlockite (green), cHAp (purple) and elastin (turquoise). h) Overview of the Mi/Ma for the mineral-rich ECM region in g. Scale bars in g and h: 10 µm. i) Demixed spectra belonging to the components identified in e and g: whitlockite (green), cHAp (purple), lipid (orange), collagen (blue), and elastin (turquoise). Identification of the mineral phases was based on the position of the v1 peak (960 cm^-1^ for cHAp and 971 cm^-1^ for whitlockite) and v4 (590 cm^-1^ for cHAp and 632 cm^-1^ for whitlockite). The lipid-rich matrix was identified mainly on the 699 cm^-1^ peak, assigned to cholesterol. Collagen and elastin were distinguished based on the desmosine, iso-desmosine and hydroxyproline peaks (see SI-2 for details).

**Fig 3.**
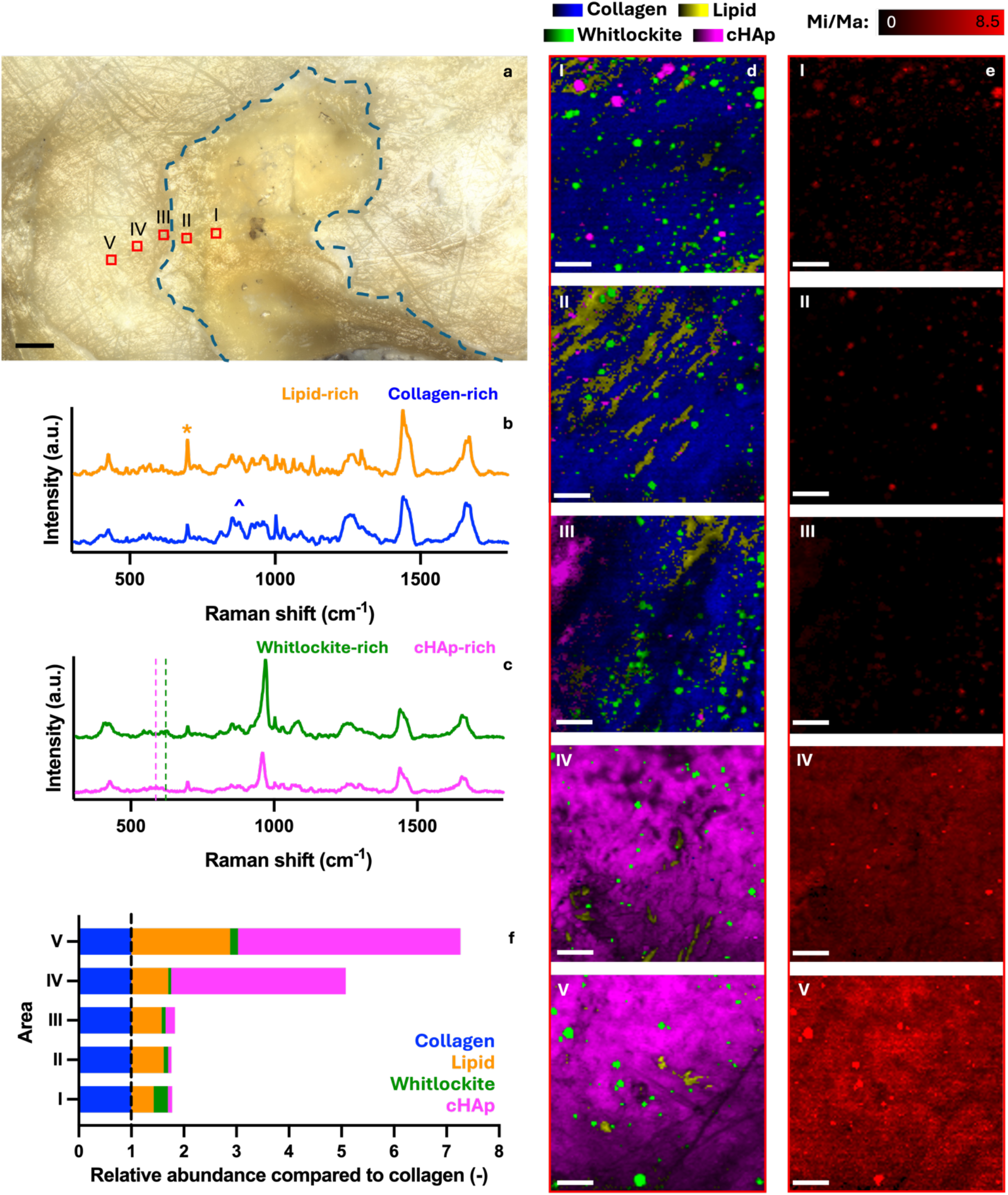
Characterization of the fully mineralized region of a calcified human aortic valve. a) Overview of the vibratomed surface of the mineralized aortic valve, with the selected regions of interest (I-V) highlighted in red. The blue dashed line indicates the border between the cHAp-rich matrix and the cHAp-poor matrix in the valve. Scale bar: 100 µm. b) Mixed component spectra of the organic contributors in area III representing the lipid-rich matrix (yellow) and the unmineralized collagen-rich matrix (blue). Lipids were identified mainly by the cholesterol (699 cm^-1^) peak and the organic matrix was identified as collagen by the presence of the (hydroxy-)proline peaks (857, 875 cm^-1^). c) Examples of mixed spectra for the cHAp-rich regions (purple) and whitlockite-rich regions (green) in area III. Identification of the mineral phase was based on the peak position of v1 (961 cm^-1^ (cHAp) vs. 974 cm^-1^ (whitlockite)) and v4 (590 cm^-1^ (cHAp) and 630 cm^-1^ (whitlockite)). d) Component images taken from regions I-V as shown in panel a. The colors in all five regions represent the major constituent in each pixel defined by the demixed spectra in Fig. 2i: collagen-rich matrix (blue), cHAp-rich matrix (purple), whitlockite-rich matrix (green), lipid-rich matrix (yellow). e) Overview of the Mi/Ma for the mineral-rich ECM regions as identified in d. Scale bars in d and e: 10 µm. f) Quantification of the contribution of the different components to the overall composition of the five areas normalized by the contribution of collagen.

#### Raman data analysis

##### General

WITec 5 Project software was used for data processing and analysis. The raw Raman spectra were baseline-corrected with a shape size of 400 for the spectra collected with the 532nm laser and 150 for the spectra collected with the 633 nm laser. Furthermore, cosmic rays were removed with the built-in algorithm.

*True Component Analysis* (TCA) was then applied to the Raman imaging data set to automatically identify the unique Raman spectra and the number of components in a sample. The automatically identified component spectra were then demixed, such they correspond to the spectra of the pure chemical components of the sample. The distribution of the chemical components was mapped across the analyzed areas, highlighting the component that contributed to the majority (> 70%) of the spectrum of each pixel. The major components identified by TCA and demixing were quantified by summing up the relative contribution of each component to all the pixels across the entire area. The relative area coverage of the components was calculated by dividing the number of pixels of the components by the total amount of pixels of the image. The contribution of each of the components was then normalized by the contribution of collagen to the overall signal of the area.

##### Quantification

As the mineral orientation proved to be random in the samples investigated, the mineral-to-matrix ratio (Mi/Ma) was determined based on the polarization-dependent v1 PO4 peak, benefitting from the high signal intensity compared to the intensities of the v2 PO4 and v4 PO4 polarization-independent signals, obtain better quantification for poorly mineralized regions.[46]

The processed data were averaged with a spatial average filter size of 1 to reduce the noise level of the data. To determine the mineral-to-matrix ratios (Mi/Ma), a field mask was drawn to only include the spectra containing a mineral peak in the 950-980 cm^-1^ region. From these spectra, the areas of the phosphate v1 peak at position 930-990 cm^-1^ for the mineral and the 1600-1720 cm^-1^ of the amide I for the organic matrix were obtained for every single pixel. The Mi/Ma was then determined by dividing the area of the phosphate v1 peak by the area of the amide I peak in every pixel.

#### Preparation of the TEM lamella

Lamellae were prepared from selected areas as described in Bertazzo et al[29]. Correlation was performed by creating visual overlays of the optical images taken using the Raman microscope and secondary electron images (Zeiss Crossbeam 550). Images were loaded in the Atlas software and coarsely correlated based on the outline of the full sample and subsequently aligned with the outline of the fracture inside the sample to achieve a fine correlation between the Raman 2D depth scan image and the TEM lift-out.

Coarse trenches were milled using a Zeiss Crossbeam 550 cryoFIB/SEM around the lamella to create access to the thick lamella using the 30 kV@30 nA FIB probe. Then, the lamella was removed from the tissue using a micro-gripper (Kleindiek, Reutlingen), attached to a half-moon copper grid (Copper lift-out grids 75964-01, Aurion, The Netherlands), and thinned. The thinning was done with a 30kV@3nA FIB probe, followed by thinning with the 30 kV@700pA and the last polishing step was done with a 30 kV@50pA FIB current to create a smooth surface on both sides of the lamella for TEM. The final lamella thickness was approximately 200 nm.

#### TEM imaging

TEM imaging was performed on a Talos F200C transmission electron microscope (Thermofischer Scientific). The half-moon copper grid was loaded into the system, and images were taken at 200kV at multiple magnifications using a Ceta-S speed camera for both imaging and diffraction measurements. The image and diffraction pattern from figure 4 a&b were made with a JEM-2100 (JEOL) equipped with a Gatan Orius CCD camera (Gatan) at 200 kV. The electron diffraction patterns were calibrated to a standard gold diffraction pattern. The patterns were indexed following Gopal et al. 1974 using the ReciPro software [45] and the azimuthal integration provided by the PASAD package [46] for Digital Micrograph (Gatan). EDX measurements were performed using a Bruker X-flash 6T_30 (Bruker).

**Fig 4.**
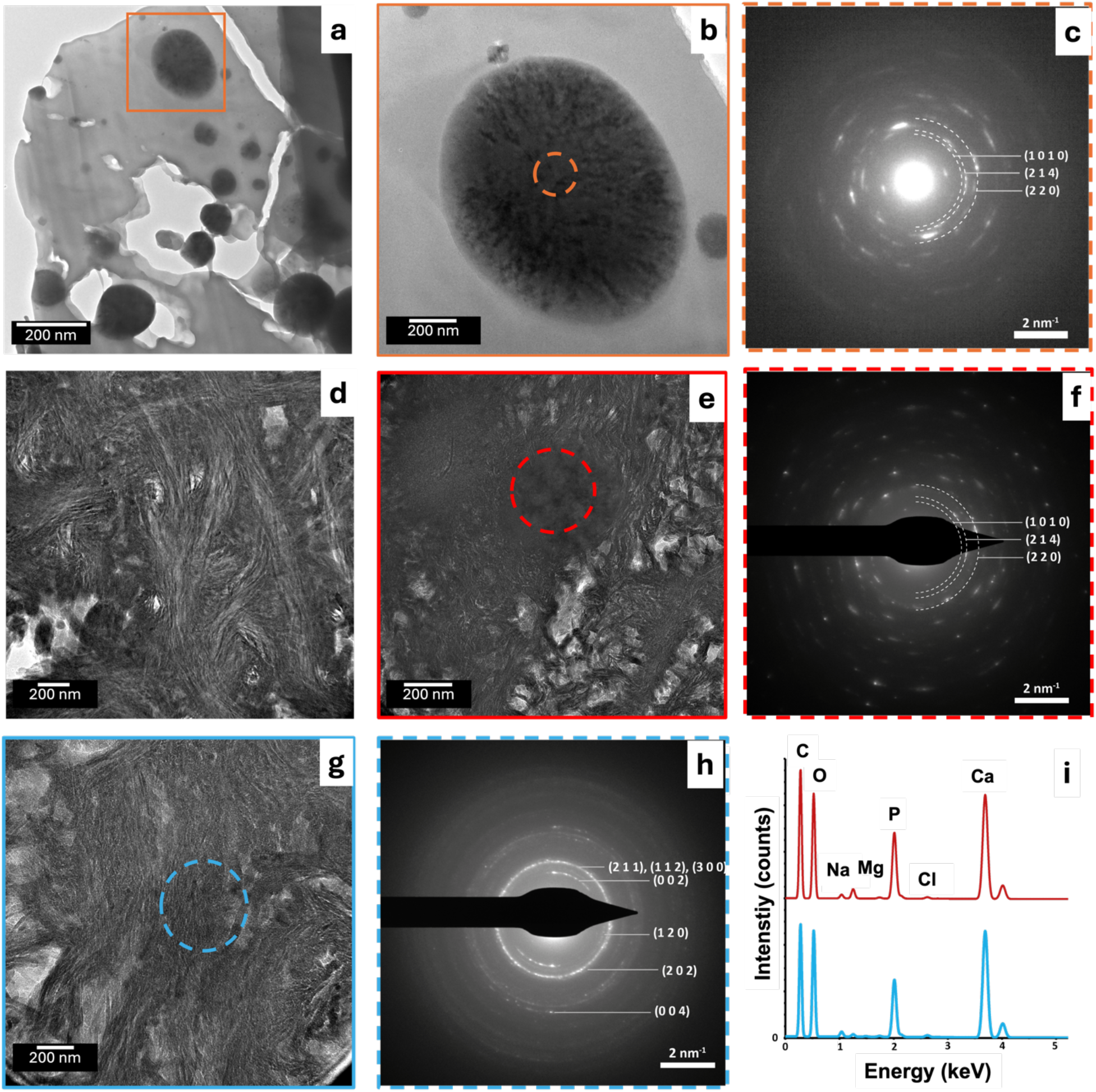
TEM analysis of a mineralized aortic valve. TEM images of lamellae created from region III (a and b) and region V (d, e and g), as indicated in Fig. 3. Lamellae were prepared using targeted FIB-SEM-milling. a) Overview of the lamella from region III, showing multiple spherical mineral particles in the matrix ranging from 100 to 1000 nm in size. b) Higher magnification image of the sphere marked by an orange box in a showing the structure of two mineral particles of 1000 nm and 150 nm in size. c) The electron diffraction pattern of the area marked in) confirms that the mineral particle in b is whitlockite. d) High magnification image of a lamella taken from the fully mineralized region V indicated in Fig. 3. shows highly directional mineral platelets following the organization of the collagen fibers. e) High magnification image of a whitlockite crystal embedded in the mineralized collagen matrix, as confirmed by f the electron diffraction pattern taken from the red circle, which is consistent with whitlockite. g) High magnification image of the same lamella showing the mineralization pattern in the valve. h) Electron diffraction pattern taken in the blue circle in g), consistent with cHAp crystals oriented along the long axis of collagen fibrils, similar to those found in bone. i) EDX spectra taken from the hydroxyapatite and whitlockite (red and blue circles in e and g, respectively). The whitlockite shows a clear increase in the Mg signal and a reduction in the Ca/P ratio.

## Results

### Mapping ECM modifications in AS

Multiple 4 µm thick sections of a freshly dissected bicuspid aortic valve of the 74-year-old male patient (Sample M-108) were cut with a microtome from a mildly mineralized region of the heart valve, with consecutive slices being selected for histology and Raman microscopy (Fig. 2). Masson’s Trichrome staining of the histological section creates a good baseline for the localized assessment of the extend CAVD had progressed in the tissue and can indicate specific areas for further in-depth investigation. The histology shows the presence of the collagenous fibrosa and the elastin-rich ventricular layers (Fig. 2a, blue and light pink, respectively, and SI-1). It further showed that part of the mineral deposit had broken out (highlighted by *) and was lost during sample preparation. Based on the correlation with the histological section, two regions in the collagen-rich fibrosa layer - which is known to be the first layer to display mineralization[47] - were selected for Raman microscopic analysis (Fig. 2b, SI-1):[48] one having no visual mineral deposits (Fig. 2a-b - blue box, 2c), and one close to the visible mineralized region (Fig. 2a-b - red box, 2d).

In the region without visual mineral deposits (Fig. 2c, 2e) true component analysis (TCA) of the Raman spectroscopy identified the main spectral component to be collagen, as indicated by the presence of the hydroxyproline (878 cm^-1^), amide III (1242 cm^-1^), ∼CH2 (1450 cm^-1^) and amide I (1666 cm^-1^) peaks (Fig. 2e, blue; Fig. 2i, blue spectrum). Cholesterol-dominated lipid deposition, as identified by its characteristic steroid ring stretch at 699 cm^-1^, as well as changes in the shape of the 1455 and 1660 cm^-1^ vibrational peaks compared to the collagen matrix, was distributed between the collagenous tissue (Fig. 2e, yellow; Fig. 2i, orange spectrum).[49, 50] Despite the oval shapes of the lipid-dominated regions in the component map that make them resemble cellular structures, both the density of the shapes and the amount of cholesterol relative to the collagen matrix suggest that the cholesterol-dominated lipid fraction is extracellular deposit. Although visually no mineral deposits were observed, surprisingly, the third spectral component showed a significant presence of whitlockite, a magnesium-substituted calcium phosphate mineral, with peaks at 972 cm^-1^ (v1 PO4) at 630 cm^-1^ (v4 PO4) vibration (Fig. 2e, green; Fig. 2i, green spectrum).[51, 52] Whitlockite has been demonstrated in mitral valve calcification,[53, 54] in other pathologically calcified tissues,[55-57] as well as in a recent laboratory model,[58] but not in human aortic valves. Here it was present in the form of micro-crystals with dimensions in the range of 1-3 µm associated with the collagenous matrix (Fig. 2e, green). The fourth minor spectral component indicated that a small amount of the collagen matrix had been mineralized with cHAp, suggesting that in this region, matrix calcification was still in its very early stages (Fig. 2e purple; Fig. 2i, purple spectrum). The local degree of mineralization was mapped, determining the Mi/Ma based on the combined intensities of the cHAp (960 cm^-1^) and whitlockite-specific (972 cm^-1^) signals, and the matrix (1660 cm^-1^) signal in each pixel. The Mi/Ma varied strongly, from a small amount of mineral associated with the collagen matrix (Fig. 2f, black to dark red color) to almost pure mineral precipitation (Fig. 2f, bright red spots). The cHAp particles showed similar Mi/Ma as the whitlockite particles (SI-2).

In the second region, which is close to a visibly mineralized region (Fig. 2d), Raman microscopy showed a significantly different matrix composition compared to the first region, specifically pointing to elastin as a significant additional organic ECM component, identified by the desmosine and iso-desmosine peaks at 530 cm^-1^ and 1103 cm^-1^ (Fig. 2g, turquoise; Fig. 2i, turquoise spectrum; SI-3). Interestingly, a comparison of the elastin spectrum of this region with a spectrum of isolated elastin and elastin of an unaffected valve revealed the reduction of two peaks specific for the elastin (iso-)desmosine cross-links at 530 cm^-1^ and 1103 cm^-1^. This indicates that the mineralized elastin in the selected area had been subject to degradation (SI-4).

A second main difference was that the most prominent mineral component in this region was identified as carbonated hydroxyapatite (Fig. 2g, purple), with typical peaks for the PO4 v1 and the CO3 (carbonate) vibrations at 960 cm^-1^ and 1072 cm^-1^, respectively (Fig. 2i, purple spectrum), while whitlockite microcrystals were only a minor component (Fig. 2g, green). As in Fig. 2e, lipid is dispersed between the other components of the tissue (Fig. 2g, yellow). In contrast to the first region, the mineralized region showed very high mineral to matrix ratios (Fig. 2h), pointing to the deposition of close-to-pure mineral.

### Spatially resolved quantification of the calcified ECM

To investigate also the more dense, heavily mineralized region of the aortic valve, the leaflet obtained from a 74-year-old female patient (M-178) was cut transversely using a vibratome, exposing a clean and flat surface without risking breakout of the mineral deposit (Fig. 3a).

The border of the calcified region could be discerned, and different regions for investigation were indicated ∼100 µm apart on a line perpendicular to this border, covering a trajectory of unmineralized and fully mineralized parts within the sample (Fig. 3a, red boxes). Five surface regions (60×60 µm) were selected (I-V, Fig. 3d) with five corresponding areas for depth scans (SI-5 and SI-6).

TCA of Raman spectra recorded in these regions identified four different matrix environments (Fig. 3b,c): noncalcified collagen-rich matrix, lipid-rich collagen matrix, cHAp-rich matrix, and whitlockite-rich matrix. The collagen-rich matrix was identified based on the (hydroxy-)proline peaks at 857 cm^-1^ and 875 cm^-1^, which differentiate it from non-collagenous proteins. The matrix was labeled lipid-rich when it had a cholesterol peak (699 cm^-1^) higher in intensity than the amide III peak (1232 cm^-1^). Raman mapping of the demixed spectra (for demixed component spectra see Fig. 2i) indeed confirmed that the cHAp-dominated areas increased going from the periphery of the calcification (regions I and II) to the border (region III) towards its center (regions IV and V) (Fig. 3d-f), with Mi/Ma ratios reaching 8.5 in a few heavily calcified areas (Fig. 3e). For comparison, Mi/Ma values for mature osteonal human bone measured under the same conditions are close to 1.5.[59]

All regions showed a significant deposition of cholesterol-rich lipids, with their abundance increasing from regions of low mineralization to those of heavy mineralization (Fig. 3f). Cholesterol-rich areas were lining the collagen-rich layers in the mildly mineralized regions (I-III), where they were also found located nearby and sometimes co-localizing with the whitlockite-rich areas. In the heavily cHAp-calcified regions (IV-V), the cholesterol-rich phase showed the same layered appearance.

Deconvolution of the PO4 v1 peak revealed that at the resolution of Raman microscopy (∼ 500 nm in X-Y), the heavily mineralized regions (IV-V) additionally showed a matrix environment where cHAp and whitlockite co-exist. In the spectra of this mixed mineral phase, which accounted for 2-3% of both measured areas (Fig. 3e), the contribution of the whitlockite amounted to between 3% and 50% of the total PO4 v1 peak area (SI-7).The presence of whitlockite even at late stages of mineralization and in co-existence with heavily mineralized apatite matrix, as well as its similar abundance in the five regions of varying overall mineralization, suggests that the whitlockite does not actively promote or precede cHAp formation, but that cHAp formation is an independent event and that the cHAp mineral grows into and engulfs the whitlockite microparticles.

### Raman-guided TEM shows mineral-matrix interactions

To understand the details of the spatial relation of the mineral crystals and the organic matrix, we used transmission electron microscopy (TEM) to further investigate the ultrastructure of the moderately (region III) and heavily (region V) mineralized ECM regions. For this aim, ∼100 nm thin electron transparent TEM lamellae were prepared using a focused ion beam / scanning electron microscope (FIB/SEM), taken from the regions where the Raman depth scans (SI-6) were performed. Here, sample features observed in the optical imaging mode of the Raman microscope were overlayed with the SEM images in the FIB/SEM to guide localization in the lamella preparation process.

A lamella taken at the border of the calcified region (region III) contained round deposits of electron-dense material with diameters of 100-1000 nm randomly dispersed throughout the matrix. (Fig. 4a) Electron diffraction of the spheres showed they were crystals of whitlockite, with slightly misaligned crystalline domains as judged from the arc in the diffraction pattern (Fig 4b,c; SI-8), [60] confirming the observations in Raman microscopy (Fig. 3d).

A second lamella was prepared from a heavily mineralized area (region V) where the cHAp-rich matrix that was identified with Raman microscopy (Mi/Ma > 6) provided strong TEM contrast (Fig. 4d). Also here, we observed individually dispersed spheres of mineralized material, with sizes between 100 and 500 nm (Fig. 4e). Also here, electron diffraction showed they were crystals of whitlockite (Fig. 4f, SI-8),[60] consistent with EDX data that showed a clear Mg peak and a Ca/P ratio of 1.53 (Fig. 4i).

The presence of a cHAp-rich collagen matrix that closely engulfed the whitlockite crystals was confirmed by electron diffraction (Fig 4g,h, SI-9) and the Ca/P ratio of 1.71 found in EDX (Fig 4i). Polarized Raman microscopy,[35] did not show any preferred orientation for either the mineral crystals nor the collagen at the multi-micron level, and TEM indicated the mineralized collagen fibers were observed to bend through the field of view in random orientation, contrasting the highly organized collagen matrix in a healthy valve.[9, 47] At the nanoscopic level the mineral crystal orientation still appeared to follow the collagen fibril direction (Fig. 4d,g). The (002) reflection arc of the cHAp in the electron diffraction indeed revealed a directional organization of the cHAp minerals, with an angular dispersion of 27° (Fig. 4h). This is very similar to what is observed in the physiological mineralization of collagen in bone, indicating that this collagen matrix serves as a template for the growth of the mineral platelets.[61, 62]

## Discussion

In the above, we highlighted the power of Raman microscopy to reveal details on the structure and chemistry of extracted calcified heart valves, focusing on its potential to aid in answering essential questions regarding the development of CAVD. Due to the low number of samples investigated, we do not claim that our observations are representative of the effects of the disease on matrix composition in large groups of CAVD patients. Nevertheless, our results do reveal the type of details that Raman microscopy can resolve and the sort of conclusions that can be drawn. The combination of Raman microscopy and TEM, where we used Raman microscopy to guide FIB/SEM lift-out and TEM analysis to locations of specific interest, provided unprecedented, detailed chemical and ultrastructural information on the matrix calcifications. Importantly, in this approach, FIB sectioning safeguarded the integrity of the mineral deposits for TEM analysis, going beyond the capabilities of conventional microtomy-based strategies.

Our experiments focused on three important unresolved aspects of the development of CAVD: 1) the chemical modification of matrix macromolecules that precedes calcification, 2) the deposition of lipids and the development of the mineral phase, and 3) the similarity/dissimilarity of matrix calcification with osteogenic mineralization.

Our results demonstrate that Raman microscopy can be used to analyze the chemistry of the matrix in detail: we quantitatively identify different degrees of mineralization, different mineral types (cHAp, whitlockite) and different calcified matrix polymers (collagen, elastin). Although the literature points to collagen as the major mineralizing matrix[48], our results indicate mineralization in places is associated with elastin domains (Fig 2g). These observations align with studies that report on matrix degradation[63-65] and intermixing of different ECM layers[9, 47] at different stages of CAVD. Indeed, we reveal that with Raman microscopy also minor matrix modifications can be detected, such as the loss of cross-links associated with the mineralized elastin domains.

Previous work has reported on the deposition of cholesterol and its unsaturated esters in calcified aortic valves[66, 67] and the possible relation between cholesterol concentrations in blood and the progression of CAVD in mice.[68] We demonstrate here how Raman microscopy can be used to quantify the cholesterol deposits within the matrix. Our observation of increased concentrations of cholesterol in the areas with severe calcification suggest that - in the material examined – cholesterol deposition is not only associated with the early stages of the CAVD process but continues during the heavy calcification of the matrix.

Our data show 100-1000 nm mineral microparticles in areas with low and moderate degrees of calcification, similar to the findings of Bertazzo et al.[29] Although the EDS analysis of their micro-particles showed the presence of calcium, phosphate, and magnesium, they have been identified as apatite. [29] Another study showed the presence of deposits of similar size particles in CAVD tissue,[58] and based on their rhombohedral shape, it was suggested that these particles were whitlockite.[51, 57]. Here, we observe only spheroidal mineral deposits with similar sizes that we identify as whitlockite using a combination of Raman microscopy, TEM imaging, EDX and electron diffraction. While cHAp is the mineral component of bone and teeth produced under physiological conditions, whitlockite is a magnesium-substituted calcium phosphate mineral absent in healthy mineralized tissues. The presence of whitlockite, however, has been demonstrated in cases of pathological mineralization beyond the aortic valve,[57] for example, in mitral valve calcification,[53, 54] upon mineralization of the arteries,[67] in breast cancer calcification,[69] as well as in an *in vitro* laboratory model,[58] but not yet in human AS.

While whitlockite has been proposed to act as a precursor of cHAp deposition,[70, 71] we detect the co-existence of cHAp and whitlockite also in areas with severe cHAp-dominated calcification. This indicates that whitlockite is not a feedstock for cHAp formation but that the two mineral phases form independently, as was very recently suggested based on an *in vitro* laboratory model.[58] Moreover, in the areas with advanced calcification, we observed the close intergrown co-localization of the two phases, implying that the cHAp engulfs the whitlockite during the development of the calcification.

The cHAp mineralized collagen, at first glance, is similar to the mineralized collagen in bone. As in bone, we find these plate-like cHAps following a weaving pattern by which the collagen matrix was mineralized. Both polarized Raman and TEM show that collagen bundles have a random organization throughout the matrix, similar to what is observed in woven bone but also in specific regions of lamellar bone.[72] However, the Mi/Ma value of region V (Mi/Ma > 6) is much higher than the M/Ma ∼ 1.5, which is more typically found in bone.[59] This suggests that the mechanisms controlling the level of mineralization in the aortic valve may be very different from those occurring in bone.[73]

The whitlockite microcrystals we observed clearly do not result from osteogenic mineralization. [57] The rounded shape of the observed whitlockite crystals suggests an immaturity in the crystal formation process, which is also reflected in the slight misalignment of crystal domains indicated by the electron diffraction patterns. Still, their near monocrystalline structure points to a slow process of crystal growth driven by low degrees of supersaturation,[74] rather than to the chaotic processes associated with dystrophic mineralization resulting from apoptotic or necrotic cell death that have been described for apatite deposition in CAVD. [75]

Although the current end-point data cannot shine light on the origin of their formation, the low concentrations of Mg^2+^ present in serum suggest that these whitlockite crystals have developed over a long period of time as the consequence of a currently unknown process. As whitlockite microcrystals may well serve as triggers for the osteoblastic differentiation of VICs and the resulting deposition of cHAp mineralized collagen matrix, the origin of their formation should be an important topic of future investigations.

## Conclusion

Aortic valve stenosis is a disease that poses a massive burden on general healthcare. Currently, treatment is limited to invasive surgical replacements due to a lack of fundamental understanding of the disease progression. In this work, we show the potential of Raman microscopy to investigate the development of compositional and structural anomalies in the ECM of stenotic aortic valves.

The presented work showcases how chemical and structural mapping by Raman microscopy allows to identify and localize different matrix biopolymers as well as mineral components in both lightly and heavily mineralized regions of the valve, and in this way, can help to understand the pathways of calcification in CAVD. This is exemplified by the observation that the degradation of elastin cross-links co-localizes with the cHAp deposition and the co-existence of whitlockite microparticles and cHAp-mineralized collage in extensively mineralized areas.

Raman microscopy does not just allow for the identification of the composition of the mineralized matrix; quantification of the composition maps allows us to better define and compare different regions with those found using other mineralization processes, such as bone formation, atherosclerosis and calcification in breast cancer tissue.

This work further shows the combined application Raman and electron microscopy, a novel multi-modal approach that allows for a multi-scale description of the ECM’s chemical and structural map and its ability to shed light on the different mineral formation mechanisms underlying CAVD. Similarly, the correlation of Raman microscopy with different electron microscopic approaches will be of great benefit for investigating matrix development in different tissue types, both in health and disease.

## Data availability

Raw data will be made publicly available on the Radboud Data Repository (DOI to be generated) and will be available upon request during the transition.

## Acknowledgements

This project was supported by the European Research Council (ERC) Advanced Investigator grant (H2020-ERC-2017-ADV-788982-COLMIN) to N.S. E.MS. is supported by the Ramón y Cajal Research Program (RYC2023-045512-I) funded by MCIN/AEI/10.13039/501100011033 and FSE+, and the project PID2022-141993NA-I00 funded by MICIU/AEI/10.13039/501100011033 and ERDF/UE. R.A. was supported by the MITACS Globalink Research Award.

## Footnotes

Additional data is provided in the supplementary information file.

