## Supplementary information for "Combined Raman Microscopy and Transmission Electron Microscopy shows the co-existence of whitlockite crystals and carbonated hydroxyapatite-mineralized collagen fibrils in human calcified aortic valves"

### S1: Overview of consecutive slices for Histological and Raman microscopy

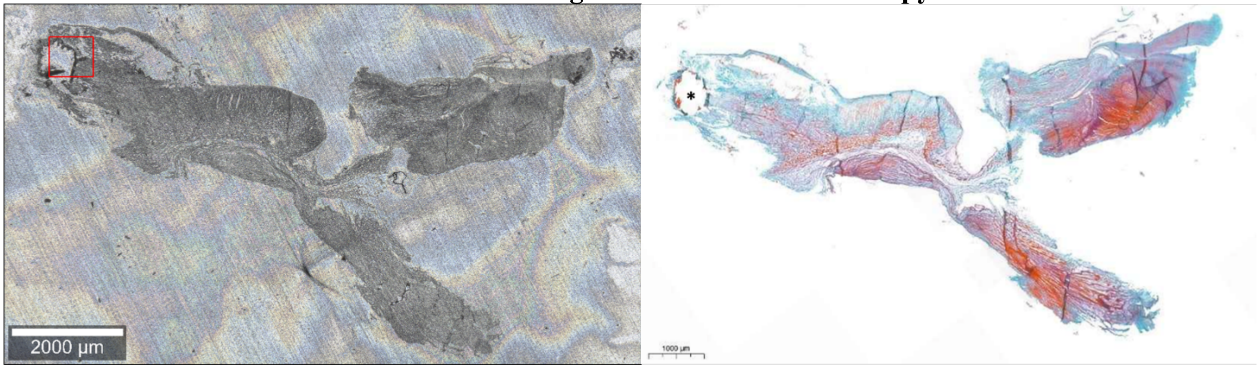


***Figure S1*** *Overview of consecutive slices.* Optical overview images of consecutive cross-sectional slices of a stenotic heart valve from the 74-year-old male patient (M-108) after cryo-sectioning. a) A slice prepared for Raman imaging placed on an aluminum substrate after cryo-sectioning. b) A slice stained with Mason's trichrome staining highlighting the presence of collagen (blue) and elastin (purple) to understand the locations of the fibrosa (top), spongiosa (middle) and ventricular (bottom) sides of the heart valve.

**S2: Mi/Ma ratio distribution of the region of valve M-108 analyzed in Fig. 2e**

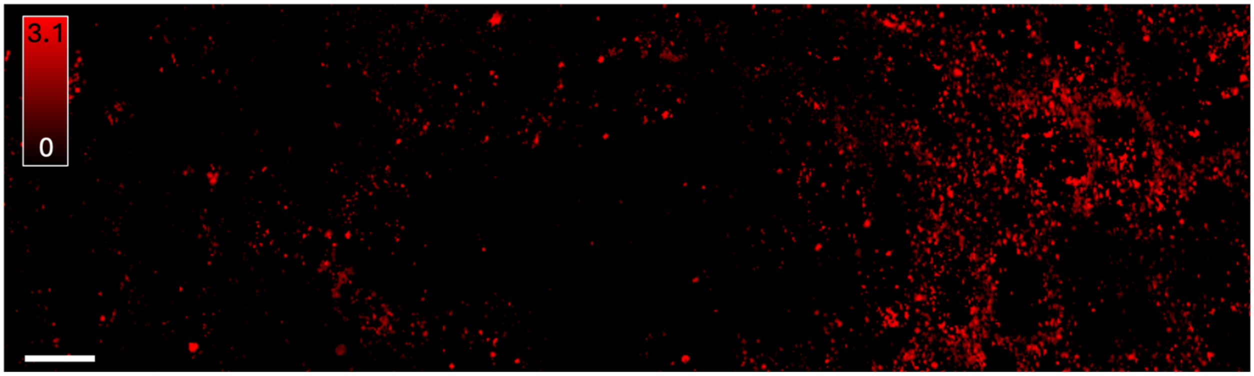


***Figure S2*** *Mi/Ma ratios of cHAp and whitlockite particles in the non-macroscopically calcified region of aortic valve M-108.* The region of the tissue shown here corresponds to the region mapped in Fig. 2e. Mi/Ma ratios of WL and cHAp in this region are comparable. Scale bar: 20 µm.

**S3: Comparison of example spectra of the map in Fig. 2g highlighting the presence of elastin and collagen**

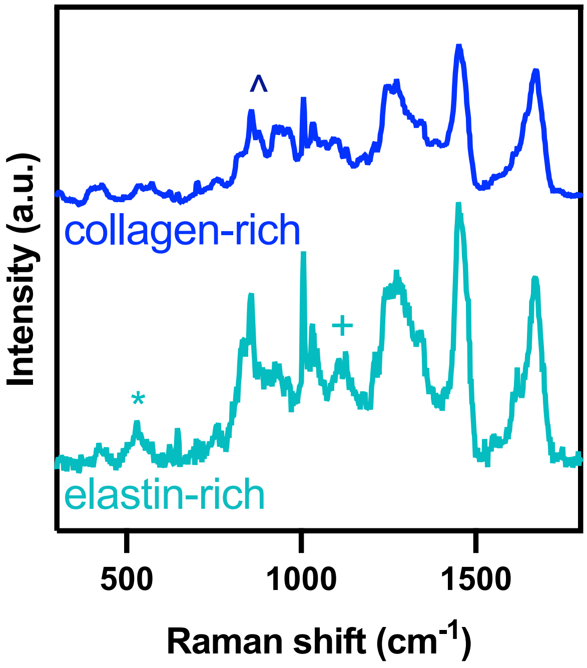


***Figure S3*** *Comparison of representative Raman spectra from a collagen-rich (blue) and an elastin-rich (turquoise) region of the calcified area of aortic valve M-108.* Collagen and elastin spectra can be distinguished by the presence of the characteristic desmosine (530 $cm^{-1}$; indicated by *) and an iso-desmosine (1103 $cm^{-1}$; indicated by +) peaks and the absence of a pronounced hydroxyproline shoulder (875 $cm-^{1}$; indicated by ^) in the elastin spectra.

**S4: Comparison of elastin spectra**

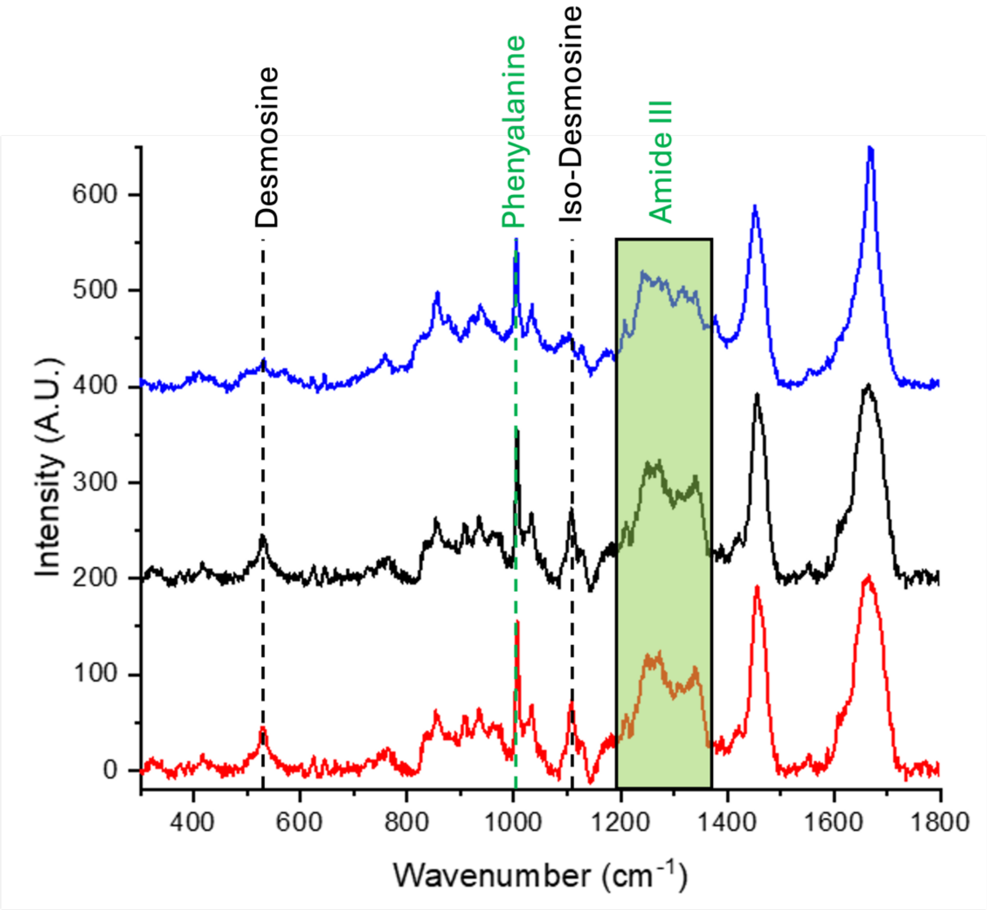


***Figure S4** Comparison of pristine and degraded elastin spectra.* Shown are representative spectra of elastin from a reference of isolated elastin (red), elastin of an unaffected heart valve (black), and elastin of a stenotic valve (blue). Comparing the isolated elastin (red) with the elastin spectra of an unaffected valve (black) we see that they are almost identical, clearly showing the features of (iso-)desmosine at 530 and 1103 $cm^{-1}$. Comparing this to the spectra of the elastin from the 74 year old male patient suffering from stenosis (M-108), we see the absence of the (iso-)desmosine signals as well as a change in the shape of the amide I (1660 $cm^{-1}$ peak) indicating degradation. The material could still be identified as elastin through the similarities in the Amide III (1200-1350 $cm^{-1}$) region as well as the spectral range from 800-1000 $cm^{-1}$.

### S5: Exact locations of area scans

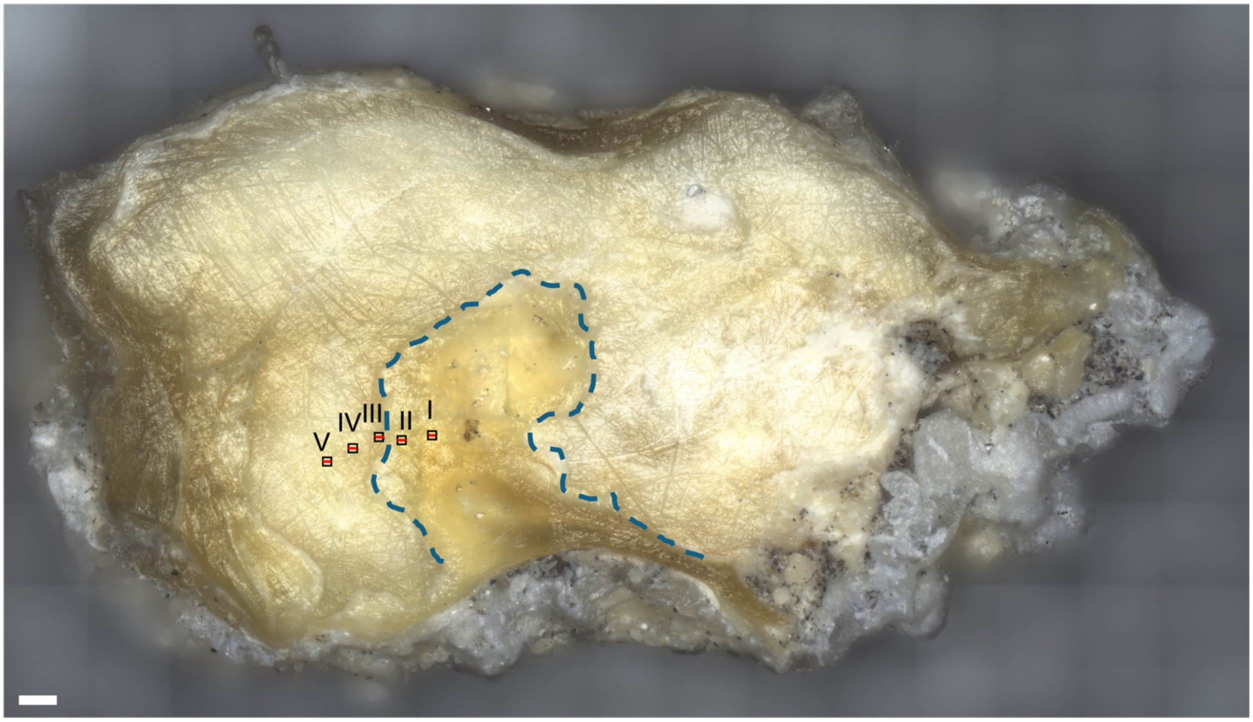


***Figure S5*** *Overview over the polished sample surface of sample M-178.* The locations of the area and depth scans are highlighted with back squares and red lines, respectively. The border between the mineralized and the non-mineralized region is indicated by the blue dashed line. Scale bar: 300 µm.

### S6: Depth scans of the regions of interest

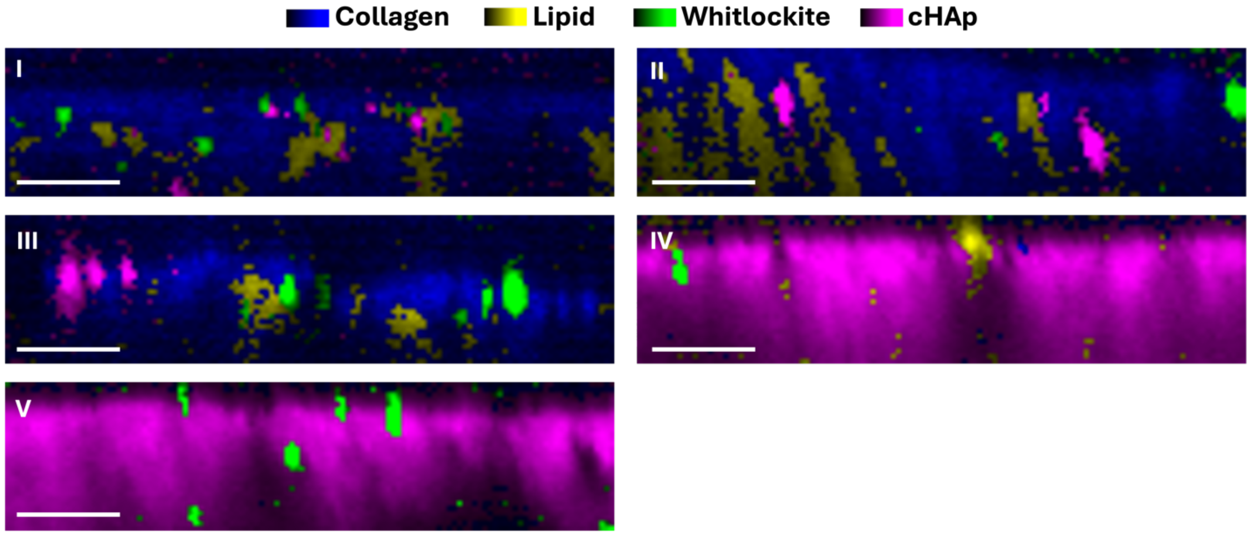


***Figure S6*** *Depths scans of the regions of interest.* Distribution of the components (spectra in Fig. 3b and 3c) in the depth scans, whose locations on the tissue and with respect to the area scans are indicated in SI-5. Scale bars: 10 µm.

### S7: Deconvolution of the whitlockite/hydroxyapatite region

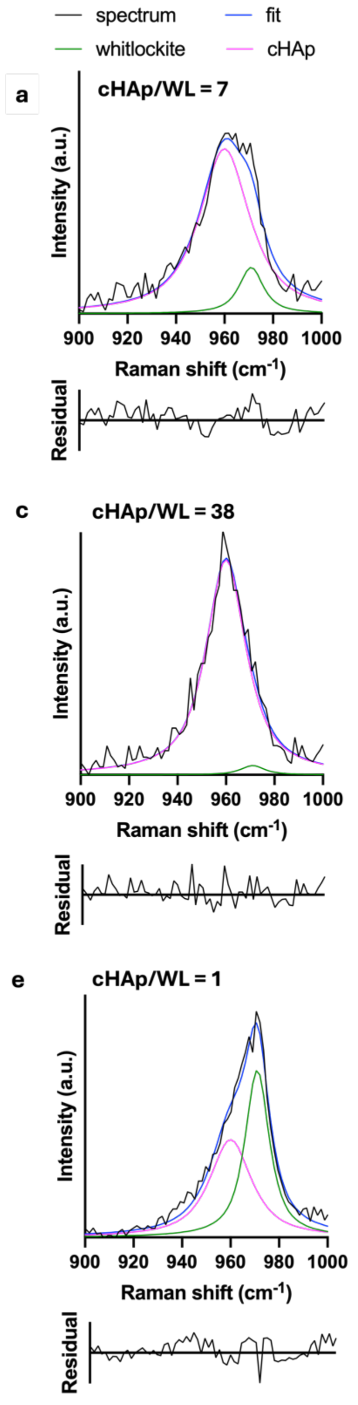


b

| Area 1 | cHAp | whitlockite |
|---|---|---|
| position | 960 $cm^{-1}$ | 971 $cm^{-1}$ |
| A | 3811 | 515 |
| HWHM | 12.5 $cm^{-1}$ | 6 $cm^{-1}$ |

d

| Area 2 | cHAp | whitlockite |
|---|---|---|
| position | 960 $cm^{-1}$ | 971 $cm^{-1}$ |
| A | 3945 | 103 |
| HWHM | 9.9 $cm^{-1}$ | 6 $cm^{-1}$ |

f

| Area 3 | cHAp | whitlockite |
|---|---|---|
| position | 960 $cm^{-1}$ | 971 $cm^{-1}$ |
| A | 2516 | 2484 |
| HWHM | 10.4 $cm^{-1}$ | 6 $cm^{-1}$ |

***Figure S7*** *Deconvolution of the whitlockite/hydroxyapatite spectral region in regions IV and V.* a, c, e: Spectral deconvolution of the single pixel spectra isolated from regions IV and V show that the v1 signature is a direct combination of hydroxyapatite and whitlockite signals with peaks at 960 $cm^{-1}$ and 971 $cm^{-1}$. The deconvolution was performed using a pseudo-Voigt model with a shape factor of 0.9 for both the whitlockite and the cHAp curves. The half width at half maximum (HWHM) of the whitlockite peak was fixed at 6 $cm^{-1}$ for better comparison between the three spectra. b, d, f: Fitting parameters (peak position (position) – fixed; area under the peak (A); HWHM – fixed for whitlockite) for the three signals were comparable for the shapes of the two major contributors. The relative areas under the deconvolved spectra where the same in both spectra where the whitlockite contribution was up to 50% to the total area under the curve.

**S8: Lattice spacing of whitlockite determined by electron diffraction (Fig. 4f) compared to X-ray diffraction references (rruff database: synthetic_database_code_amcsd 0012160 [1]).**

| Lattice plane | Experimental d-spacing Whitlockite Electron Diffraction (Å) – Fig. 4c | Experimental d-spacing Whitlockite Electron Diffraction (Å) – Fig. 4f | Reference d-spacing Whitlockite X-ray diffraction (Å) [1] |
|---|---|---|---|
| 2 2 0 | 2.58 | 2.60 | 2.59 |
| 2 1 4 | 3.19 | 3.16 | 3.189 |
| 1 0 10 | 3.45 | 3.43 | 3.43 |

[1] R. Gopal, C. Calvo, J. Ito, W. K. Sabine, "Crystal structure of synthetic Mg-whitlockite, Ca18Mg2H2(PO4)14" Canadian Journal of Chemistry, vol. 52, p. 1155-1164, 1974.

**S9: Lattice spacing of cHAp determined by electron diffraction (Fig. 4h) compared to X-ray diffraction references (rruff database: xt371_database_code_amcsd 0003642 [2]).**

| Lattice plane | Experimental Hydroxyapatite Electron Diffraction (Å) – Fig. 4h | Reference cHAp X-ray diffraction (Å) [2] |
|---|---|---|
| 0 0 2 | 3.44 | 3.44 |
| 1 2 0 | 3.09 | 3.10 |
| 2 1 1, 1 1 2 | 2.82 | 2, 79 - 2.83 |
| 2 0 2 | 2.61 | 2.64 |
| 0 0 4 | 1.72 | 1.72 |

[2] M.E. Fleet, X. Liu, P.L. King, "Ca5 P2.829 O13.686 C.26" American Mineralogist, vol. 89, p. 1422-1432, 2004.
